# Genome evolution following an ecological shift in nectar-dwelling *Acinetobacter*

**DOI:** 10.1101/2023.11.02.565365

**Authors:** Vivianna A. Sanchez, Tanya Renner, Lydia J. Baker, Tory A. Hendry

**Affiliations:** Department of Microbiology, Cornell University, Ithaca, New York, USA; Department of Entomology, The Pennsylvania State University, University Park, USA

**Author notes:** Address correspondence to Tory A. Hendry. Current address: Department of Marine Biology and Ecology, University of Miami, Florida, USA.

## Abstract

The bacterial genus *Acinetobacter* includes species found in environmental habitats like soil and water, as well as species adapted to be host-associated or pathogenic. High genetic diversity may allow for this habitat flexibility, but the specific genes underlying switches between habitats are poorly understood. One lineage of *Acinetobacter* has undergone a substantial habitat change by evolving from a presumed soil-dwelling ancestral state to thrive in floral nectar. Here we compared the genomes of floral-dwelling and pollinator-associated *Acinetobacter*, including newly described species, with genomes from relatives found in other environments to determine the genomic changes associated with this ecological shift. Following one evolutionary origin of floral nectar adaptation, nectar-dwelling *Acinetobacter* species have undergone reduction in genome size compared to relatives and have experienced dynamic gene gains and losses as they diversified. We found changes in gene content underlying carbohydrate metabolism and nitrogen metabolism, which we predict to be beneficial in nectar environments. Gene losses follow a pattern consistent with genome streamlining, whereas gains appear to result from both evolutionary divergence and horizontal gene transfer. Most notably, nectar-dwelling *Acinetobacter* acquired the ability to degrade pectin from plant pathogens and the genes underlying this ability have duplicated and are under selection within the clade. We hypothesize that this ability was a key trait for adaptation to floral nectar, as it could improve access to nutrients in the nutritionally unbalanced habitat of nectar. These results identify the genomic changes and traits coinciding with a dramatic habitat switch from soil to floral nectar.

## Introduction

The gammaproteobacteria genus *Acinetobacter* is diverse and includes (1–3). These taxa inhabit a broad range of environments, including soil and water (1–3). Some *Acinetobacter* lineages have also evolved to be host-associated or animal pathogens, with a notable example being the recently emerged human pathogen *Acinetobacter baumannii* (4). Strains in the genus are phenotypically and genetically diverse and frequently adapt to new ecological niches (5, 6). However, few direct connections have been made between specific genomic changes and ecological transitions within *Acinetobacter*, or in bacteria more broadly (7). One poorly characterized habitat transition within the genus *Acinetobacter* is adaptation for growth in floral nectar. Several *Acinetobacter* species found in floral nectar appear to be most closely related to soil-dwelling relatives (4, 8). Nectar represents a significant environmental shift compared to soil habitats, likely with different selective pressures. Genomic comparisons between *Acinetobacter* adapted to floral nectar versus other habitats can uncover how bacteria evolve to new environments and which genetic traits facilitate major ecological switches.

The high genetic diversity and genomic plasticity within *Acinetobacter* may be driven by mechanisms facilitating horizontal gene transfer (HGT), including competence for natural transformation (9–12), conjugative abilities, and prevalent mobile elements such as plasmids, prophage, and insertion sequences (8, 13, 14). Horizontally acquired genomic islands are common observed throughout the *Acinetobacter* genus (15, 16) and can contain genes conferring beneficial phenotypes like antibiotic resistance and plasmid mobilization (17, 18). HGT is a source of evolutionary novelty in bacteria (19), but other sources of genetic diversity can also be important, such as error-prone polymerases in *A. baumannii* (4, 8, 20), or gene duplication followed by divergence (21). Gene duplication can also potentially lead to increased gene expression, allowing for enhanced nutrient acquisition, temperature stress tolerance, and overall resistance to antibiotics and pesticides (22), but it is unclear how broadly important this mechanism is for bacterial adaptation.

Genetic novelty in bacteria can allow for the evolution of new traits and subsequent exploitation of new niches (23–25). In some cases, specific genetic elements have been linked to habitat-specific fitness. For instance, *A. baumannii* has resistance islands with mobile genetic elements and antibiotic resistant genes, allowing for persistence in hospital settings (6, 26).

Antibiotic resistance is a common example of a novel trait resulting from a specific environmental selective pressure because it is easily observable and important in well studied pathogen systems. Other traits that are relevant in natural systems have occasionally been connected to ecological changes in bacteria (27, 28), but such connections can be difficult to infer. In other cases, traits that are linked to success in a specific environment may be known, but not their genetic basis. For instance, the ability to access nutrients from pollen is a unique and potentially beneficial trait in floral nectar-dwelling *Acinetobacter*, but how this trait was gained is unknown (29).

Floral nectar is a nutritional reward produced by flowers to attract pollinating animals. It is high in carbohydrates; the sugar content in floral nectar can reach 90% of nectar dry weight (30, 31) and it is a resource for microbes as well (32, 33). However, floral nectar habitats create several stresses for microbes, including limitation of nutrients other than sugar (34–38).

Although nectar contains amino acids and lipids (39), it can contain limiting amounts of nitrogen for some microbes (40, 41). These factors make nectar a selective environment and can lead to strong priority effects where early arriving microbes prevent subsequent colonization of flowers (40, 42).

Culture dependent and independent methods have revealed diverse microbes that thrive in these conditions (43–49). The genus *Acinetobacter* makes up a high proportion of bacterial taxa in floral nectar and is prevalent and readily cultured from nectar environments (44, 50).

*Acinetobacter* is also frequently found associated with floral visitors. For example, *Acinetobacter apis* was isolated from the gut of the western honey bee, *Apis mellifera*, and bee pollen provisions and nests sometimes include *Acinetobacter* (51–54). However, it is unknown whether *Acinetobacter* found with pollinators are nectar-dwelling species being dispersed by floral visitors, or if they are specific associates of pollinators. For ease here, we refer to strains isolated from both nectar and floral visitors as nectar-dwelling strains.

Previous phylogenomic analysis of *Acinetobacter* isolates from nectar and bees found that they were closely related to soil-dwelling species (50). This previous work suggested one evolutionary origin of nectar-dwelling/bee association but did not assess evolutionary patterns within this lineage. We do so here, with and additional 15 genomes from nectar-dwelling isolates. These include the genomes of three previously described species, *A. apis*, *A. boissieri*, and *A. nectaris* (45, 51), newly sequenced *A. nectaris* isolates, and newly sequenced genomes of three recently described species, *A. pollinis*, *A. rathckeae*, and *A. baretiae* (50). For comparison, we included genomes from *A. brisouii*, which is isolated from soil and water and was previously found to be the closest relative of *A. nectaris*, as well as those from eight other environmental *Acinetobacter* species. We hypothesized that the switch to floral nectar from soil would drastically change the selective pressures experienced by this *Acinetobacter* lineage, leading to changes in gene content. We used comparative genomics to understand which genes may become redundant or beneficial for bacteria in floral nectar, and to identify acquired genes that may have facilitated this environmental switch.

## Results and Discussion

### Phylogeny and genome characteristics of nectar-dwelling Acinetobacter

To understand the evolutionary history of nectar-dwelling *Acinetobacter*, we constructed a phylogenomic tree using genomes of *Acinetobacter* isolates from floral nectar and floral visitors. The isolates collected from floral nectar and pollinators form a clade, with a bootstrap support of 100, separate from soil, water, and animal dwelling *Acinetobacter* species (Fig. 1).

**Figure 1.**
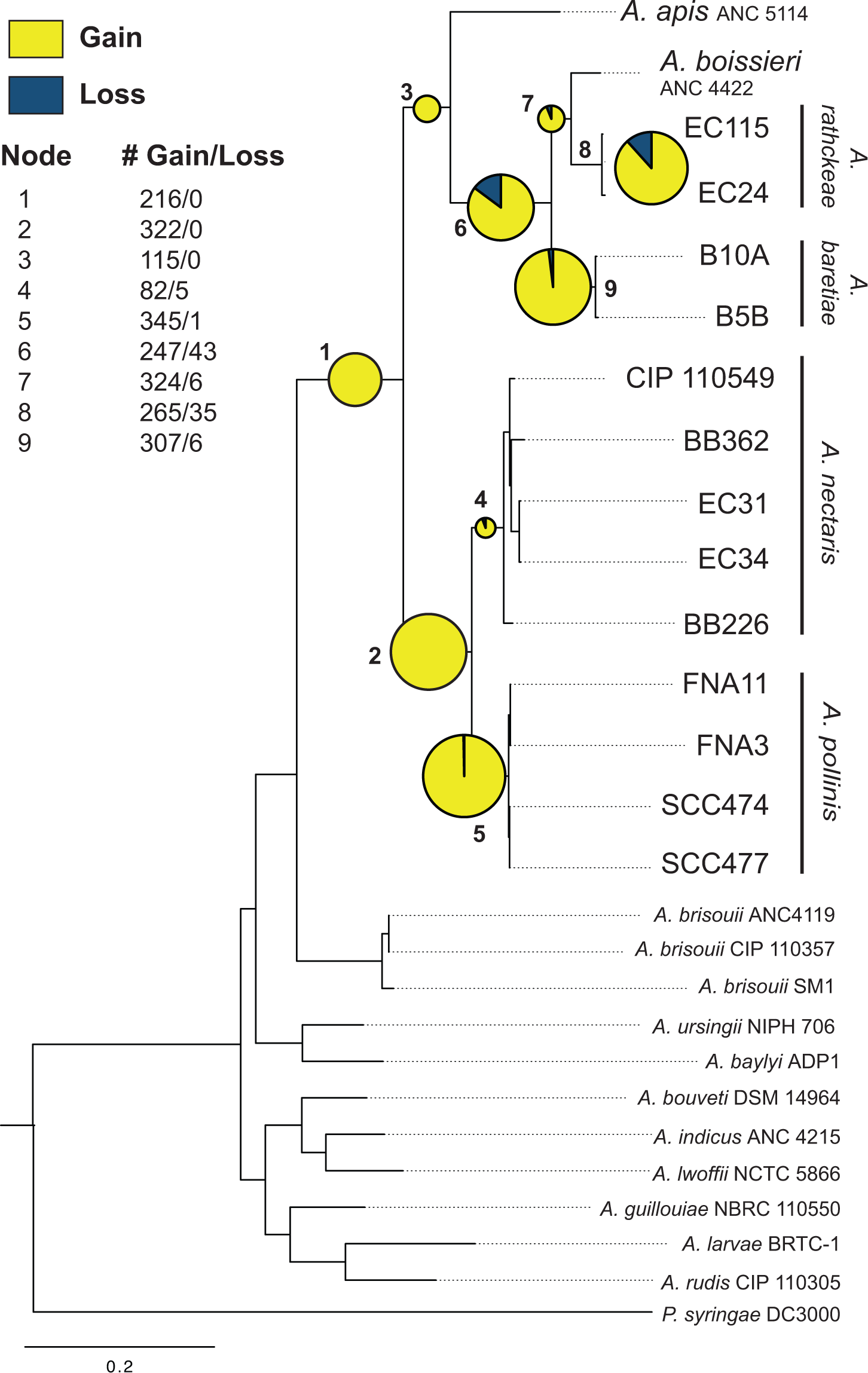
*Acinetobacter* species maximum likelihood phylogenomic tree based on 399 conserved protein sequences. All nodes have a bootstrap value of 100. Ancestral state reconstruction of ortholog gains and losses at nodes within the nectar-dwelling clade are shown. Nodes are labelled numerically and a pie chart at each node shows the proportion of ortholog gains (yellow) versus losses (blue). Pie size is scaled by total number of gain and loss events.

This confirms that there is one known evolutionary origin of a nectar-dwelling within *Acinetobacter* and that this group evolved from a presumed soil-dwelling ancestor. The six species in the nectar clade appear to not be isolated from environments outside of floral nectar or pollinators, based on 16S rRNA sequence comparisons to GenBank databases (50). Multiple of these species, *A. nectaris*, *A. boissieri*, and *A. pollinis*, are abundant and common in floral nectar from locations worldwide, and our isolates came from both North America and Europe (45, 50, 55, 56). This suggests that the clade is specialized for growth in floral nectar and/or associated with pollinators and is widely found in these habitats (44, 57, 58).

We used genomic comparisons between nectar-dwelling *Acinetobacter* and relatives living in distinct environments to uncover genomic patterns associated with nectar specialization. Relative to species found in other environments, species in the nectar-specialized *Acinetobacter* clade have smaller genomes and lower numbers of protein coding genes (Table 1). There is a significant difference between genome sizes within the nectar-specialist clade, 2.38-2.75 Mb, and the environmental clade, 2.90-4.88 Mb (p-value=0.00023). Across genomes of nectar-dwelling strains vs. environmental strains, nectar specialists have 243-977 fewer protein coding genes, a 10-30% reduction in proteins. Genomic reduction can occur for various reasons.

**Table 1.** Genome characteristics of nectar-dwelling Acinetobacter strains and comparison environmental strains. Type strains are designated.

| Strain | Accession | Pseudo-<br>genes | Genome<br>Size<br>(Mb) | G+C<br>Content<br>(%) | Protein<br>Coding<br>Genes | Proteins<br>assigned<br>to COGs |
| --- | --- | --- | --- | --- | --- | --- |
| *A. apis* ANC 5114<sup>T</sup> | FZLN000000000 | 125 | 2.41 | 38.3 | 2232 | 1985 |
| *A. boissieri* ANC 4422<sup>T</sup> | NZ_FMYL000000000 | 182 | 2.69 | 38.0 | 2554 | 2378 |
| *A. rathckeae* EC115 | VTDP000000000 | 192 | 2.62 | 39.2 | 2438 | 2388 |
| *A. rathckeae* EC24 | VTDO000000000 | 214 | 2.75 | 39.3 | 2565 | 2531 |
| *A. baretiae* B10A<sup>T</sup> | VTDM000000000 | 234 | 2.59 | 37.5 | 2531 | 2402 |
| *A. baretiae* B5B | VTDL000000000 | 253 | 2.70 | 37.5 | 2629 | 2486 |
| *A. nectaris* CIP 110549<sup>T</sup> | NZ_AYER000000000 | 189 | 2.67 | 36.7 | 2569 | 2458 |
| *A. nectaris* BB362 | JAEQDL000000000 | 249 | 2.38 | 36.9 | 2269 | 2173 |
| *A. nectaris* EC31 | JAERJC000000000 | 200 | 2.42 | 36.7 | 2312 | 2303 |
| *A. nectaris* EC34 | JAERJB000000000 | 212 | 2.39 | 36.8 | 2288 | 2284 |
| *A. nectaris* BB226 | JAEQDM000000000 | 255 | 2.58 | 36.7 | 2421 | 2312 |
| *A. pollinis* FNA11 | VTDS000000000 | 245 | 2.67 | 36.6 | 2621 | 2599 |
| *A. pollinis* FNA3 | VTDT000000000 | 248 | 2.67 | 36.6 | 2615 | 2575 |
| *A. pollinis* SCC474 | VTDR000000000 | 294 | 2.75 | 36.7 | 2623 | 2605 |
| *A. pollinis* SCC477<sup>T</sup> | VTDQ000000000 | 262 | 2.75 | 36.7 | 2634 | 2609 |
| *A. brisouii* ANC4119 | NZ_APPR000000000 | 187 | 3.11 | 41.7 | 3036 | 3021 |
| *A. brisouii* CIP 110357<br>= DSM18516 | NZ_AYEU000000000 | 184 | 3.09 | 41.6 | 3025 | 3003 |
| *A. brisouii* SM1 | NZ_JZRE000000000 | 189 | 2.99 | 41.8 | 2913 | 2651 |
| *A. baylyi* ADP1 |  | 222 | 3.6 | 40.4 | 3376 | 2897 |
| *A. ursingii* NIPH 706 | NZ_APQB000000000 | 234 | 3.49 | 40 | 3468 | 2920 |
| *A. bouveti* DSM 14964<br>= CIP 107468 | NZ_APQD000000000 | 198 | 3.37 | 40 | 3468 | 2657 |
| *A. indicus* ANC 4215 | ATGH01000001 | 238 | 3.17 | 45.4 | 3064 | 2632 |
| *A. lwoffii* NCTC 5866 =<br>CIP 64.10 = NIPH 512 | NZ_AYHO000000000 | 268 | 3.21 | 43 | 3107 | 2635 |
| *A. guillouiae* NBRC<br>110550 | NZ_APPJ000000000 | 258 | 4.88 | 38.1 | 4648 | 3362 |
| *A. larvae* BRTC-1 | NZ_CP016895 | 283 | 3.74 | 41.6 | 3453 | 2598 |
| *A. rudis* CIP110305 | ATGI000000000 | 241 | 4 | 39.1 | 3748 | 2849 |

Genome streamlining is common for species living in stable, nutrient poor conditions such as some soil and marine habitats (59–62), and is thought to be driven by selection and facilitated when bacteria have large effective population sizes (59, 63–65). Some environmental stresses may also promote genome streamlining due to selection (66, 67). Gene loss can also be degenerative and result from genetic drift, with extreme examples occurring in bacteria that are host-restricted and experience frequent population bottlenecks (68–71).

Although nectar-dwelling bacteria may experience population bottlenecks due to the transient nature of the floral environment, we do not find strong evidence for genetic drift as is seen in host-restricted bacteria; such species often show high evolutionary rates, high rates of pseudogenes, and low genome GC content (70, 72). Evolutionary rate tests in PAML found no support for a faster rate in nectar-dwelling *Acinetobacter*, compared to the null hypothesis of a global clock across the nectar and environmental *Acinetobacter* phylogeny (69). Similarly, we did not find evidence of genomic degeneration in the form of pseudogenes, as the number of pseudogenes detected in nectar-dwelling *Acinetobacter* ranged from 125-294, while environmental *Acinetobacter* had a similar range of 187-283 and other *Acinetobacter* species fall within this range as well (Table 1) (73, 74). The only pattern found that is potentially supportive of genetic drift is that nectar isolates have slightly lower percent GC compositions relative to soil-dwelling species (Table 1). We therefore hypothesize that the habitat switch from soil to nectar altered selective pressures on gene content in nectar dwelling species and allowed for the loss of some functions through genome streamlining. To further investigate this, we sought to define the gene content and functional capacities of nectar-dwelling species compared to soil-dwelling relatives.

### Gene content evolution with the switch to nectar

To determine content of predicted proteins among *Acinetobacter* clades, we performed an ortholog clustering analysis to identify shared orthologs, recent paralogs, and unique genes (75) (Table S1). This analysis resulted in 7,334 orthologs in total identified across all genomes, 1,076 of which were present across all genomes. Environmental strains had 5,558 orthologs in total, with 2,921 orthologs only found in environmental species, and 111 environmental specific orthologs shared across these species. The nectar-dwelling clade had 4,413 total orthologs, with 1,776 orthologs only found in nectar-dwelling species, and 53 of these nectar specific orthologs found in all nectar clade strains.

To trace gene gain and loss events within the nectar-specialist clade, we performed a maximum likelihood ancestral state reconstruction analysis. Overall, there have been dynamic gene gains across the evolution of the group. Substantial ortholog gain events occurred at the ancestral node for the nectar-dwelling clade, the ancestral nodes to the species *A. pollinis, A. rathckeae,* and *A. baretiae,* and at additional ancestral nodes (Fig.1, nodes 1, 2, 5, 6, 8 and 9). Gene gains were significantly higher than gene losses throughout the nectar-dwelling clade, with gains of 209-345 orthologs at tips and nodes (Fig. 1, Fig. S1). Numbers of gene loss events generally increased closer to the tips of the tree (Fig. 1, Fig. S1), suggesting that genome streamlining may be a relatively recent process in the clade. Given that the nectar-dwelling genomes are smaller in total size than the genomes of relatives (Table 1), these patterns suggests that some gene gains identified in this analysis are the result of divergence leading to novel orthologs rather than horizontal acquisition of new genes alone. Selective pressures from environmental changes can increase processes like mutation rate, leading to diversification in orthologs (76). Here, adaptation to the novel nectar environment may have led to increased gene divergence.

Gene gains and losses occurred across diverse functional categories in the nectar- dwelling clade (Fig. 2). We investigated specific changes in gene content in functional categories, with the largest differences between nectar-dwelling species and environmental relatives occurring in the categories of metabolism of aromatic compounds, nitrogen metabolism, and carbohydrate metabolism.

**Figure 2:**
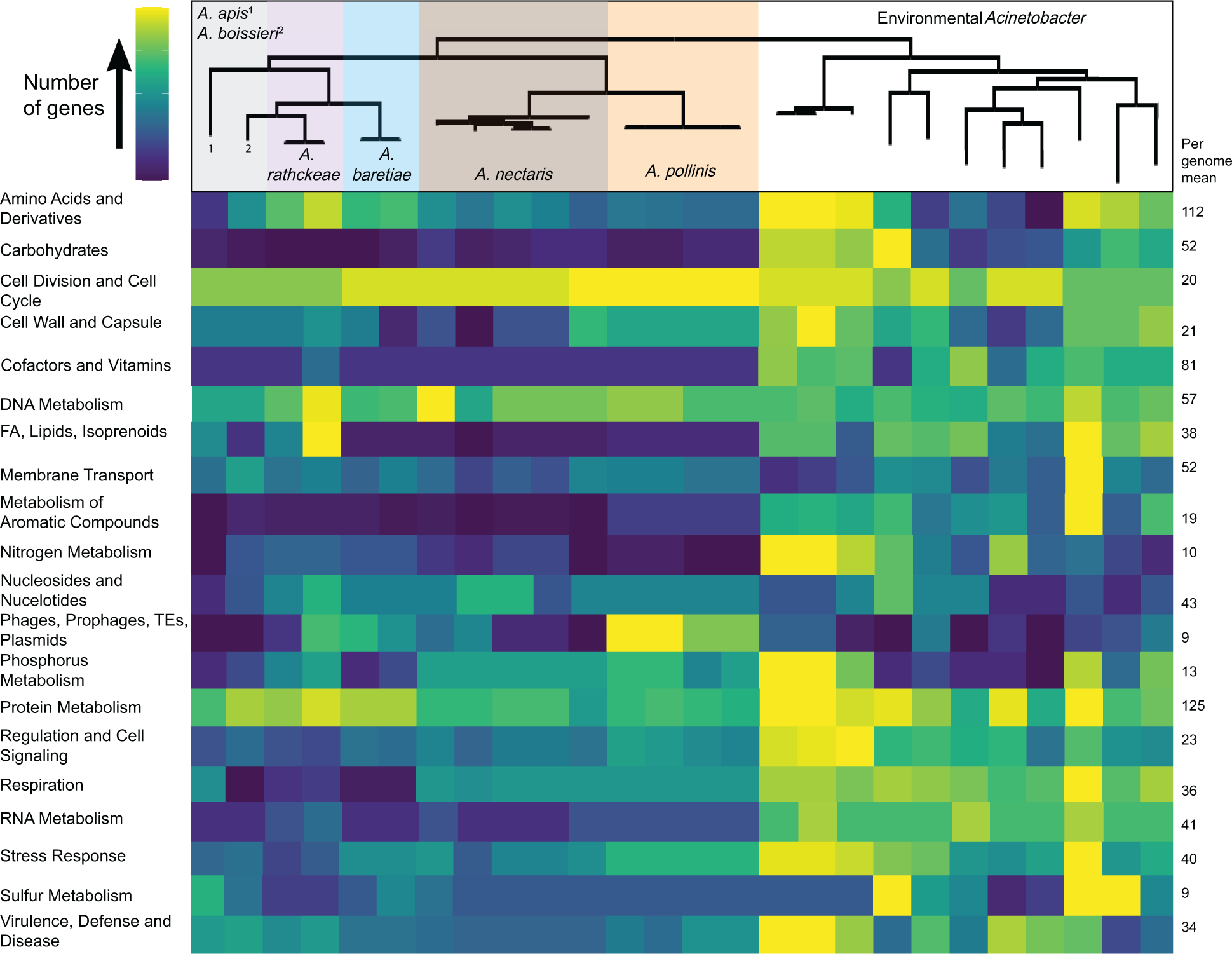
Heatmap showing number of orthologs across functional categories from nectar-dwelling and environmental *Acinetobacter* strains. Category assignments are based on RAST annotations (FA = fatty acid metabolism, TE = transposable elements), and a phylogenomic species tree (Figure 1) is shown. Each row is colored independently based on the variance of gene content within the category. A color scale with yellow indicating the highest number of genes and deep blue indicating the lowest number of genes is used. Genes of unknown function or categories with <6 mean orthologs per genome are excluded. The mean number of orthologs within a genome for each category is given.

The number of genes involved in metabolism of aromatic compounds (Fig. 2, Fig. S2A) varied between nectar and environmental strains. Although the difference in total genes was not significant, 48 out of 67 genes in this category were present in at least one environmental *Acinetobacter* species, but completely absent in the nectar clade (Table S1). All but one of the remaining genes were present in both clades, but rarely common across all nectar-dwelling strains. Several of the genes for aromatic compound metabolism that are present in nectar- dwelling *Acinetobacter* strains are involved in benzoate metabolism, an aromatic compound that has been shown to be released by plants (77). The decrease in genes in this category suggests that nectar-dwelling *Acinetobacter* encounter a limited diversity of aromatic compounds compared to species in other environments.

Gene content differences in nectar-dwelling strains compared to environmental relatives also suggest that shifts in nitrogen and amino acid metabolism strategies accompanied the shift to nectar dwelling. Genes involved in amino acid metabolism were significantly higher in nectar- dwelling strains (p-value=0.003). This difference was mainly due to an excess of genes for the transport of certain nitrogen sources, including putrescine, proline, glycine, serine, alanine, arginine, ornithine, methionine, leucine, glutamate, and aspartate (Table S1). In contrast, nitrogen metabolism genes were reduced by about half in nectar dwelling *Acinetobacter* compared to environmental species, although the difference was not significant (Table S1). Specifically, nectar-dwelling species genomes are missing glutamine synthases, glutamate synthases and ammonium transporters. Many of these are genes for which multiple orthologs are present in environmental *Acinetobacter* and only one has been retained in nectar-dwelling strains. Floral nectar is low in nitrogen relative to carbon (31, 78), and the ability to assimilate nitrogen sources has been linked to competition and growth in floral nectar in both yeasts (79) and *Acinetobacter* (80, 81). A shift towards more transport systems for nitrogen sources could be driven by selection for nitrogen scavenging.

Genes involved in carbohydrate metabolism showed an overall decrease in nectar-dwelling strains (Fig.2), however this overall decrease was not statistically significant (p-value=0.089). The loss in carbohydrate genes was driven by the subcategories of central carbohydrate metabolism and significantly by genes in the category of organic acid metabolism, (p-value=2.2e-16) (Fig. S2B). Other subcategories, notably monosaccharide metabolism, showed a significant gain in genes (p-value=3.022e-16) (Fig. S2B). Compared to environmental habitats, floral nectar provides an abundance of simple carbohydrates, including fructose, sucrose, and glucose (78, 82). Many of the nectar-dwelling species can assimilate fructose, and some can assimilate glucose and sucrose (50, 80). In comparison, non-nectar dwelling species such as *A. baylyi*, are often unable to utilize fructose, sucrose, or glucose as sole carbon sources, but can metabolize other diverse carbon sources (83–85). In support of these observations, we found that nectar-dwelling species tend to have more genes for metabolizing and transporting simple sugars (Table S1). For example, all nectar dwelling species have the gene coding for 1- phosphofructokinase (EC 2.7.1.56), which is part of the fructose and mannose metabolism pathway, whereas the genomes of environmental strains were less likely to have this gene. Nectar-dwelling strain genomes were also more likely to have the gene encoding xylanase, which is responsible for the breakdown of the common plant polysaccharide xylan to xylose (86, 87). These genes may be more beneficial for nectar-dwellers than soil-dwelling *Acinetobacter*.

In keeping with overall changes in carbohydrate metabolism in nectar, we found that phosphotransferase system (PTS) genes specific to nectar sugars are more common in nectar-dwelling strains compared to environmental strains (Table 2). PTS genes are a common method for bacteria to transport sugars into cells via a phosphorylation cascade (88, 89). PTS can also be involved in sensing and regulation of physiological processes related to sugar, such as carbohydrate active enzymes, chemotaxis, and biofilm formation (90). These multicomponent systems are specific to distinct molecules including fructose, mannitol, sucrose, and glucose. The sucrose-specific PTS enzyme complex (EIIABC) is present within *A. apis*, *A. boisseri*, *A. rathckeae*, and *A. baretiae* strains, but absent from all environmental strains, as well as *A. nectaris* and *A. pollinis*. Fructose-specific EIIABC complexes were found in all nectar-dwelling strains and only three environmental species, *A. lwoffii*, *A. baylyii*, and *A. brisouii*. This pattern suggests that PTS enzymes may have been beneficial for making the ecological switch to high sugar environments. Overall, these findings show a shift in carbohydrate metabolism in keeping with growth in nectar, although we found mainly subtle differences in gene content.

**Table 2.**
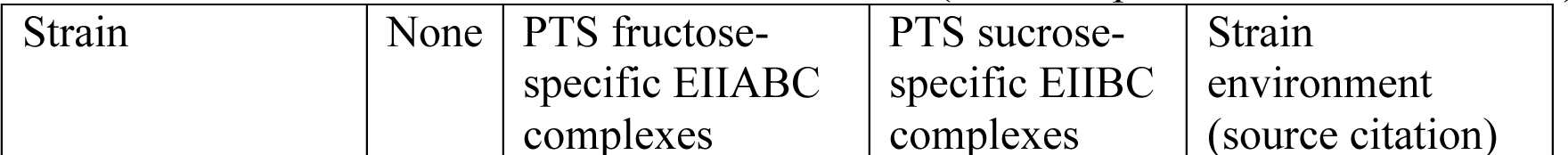
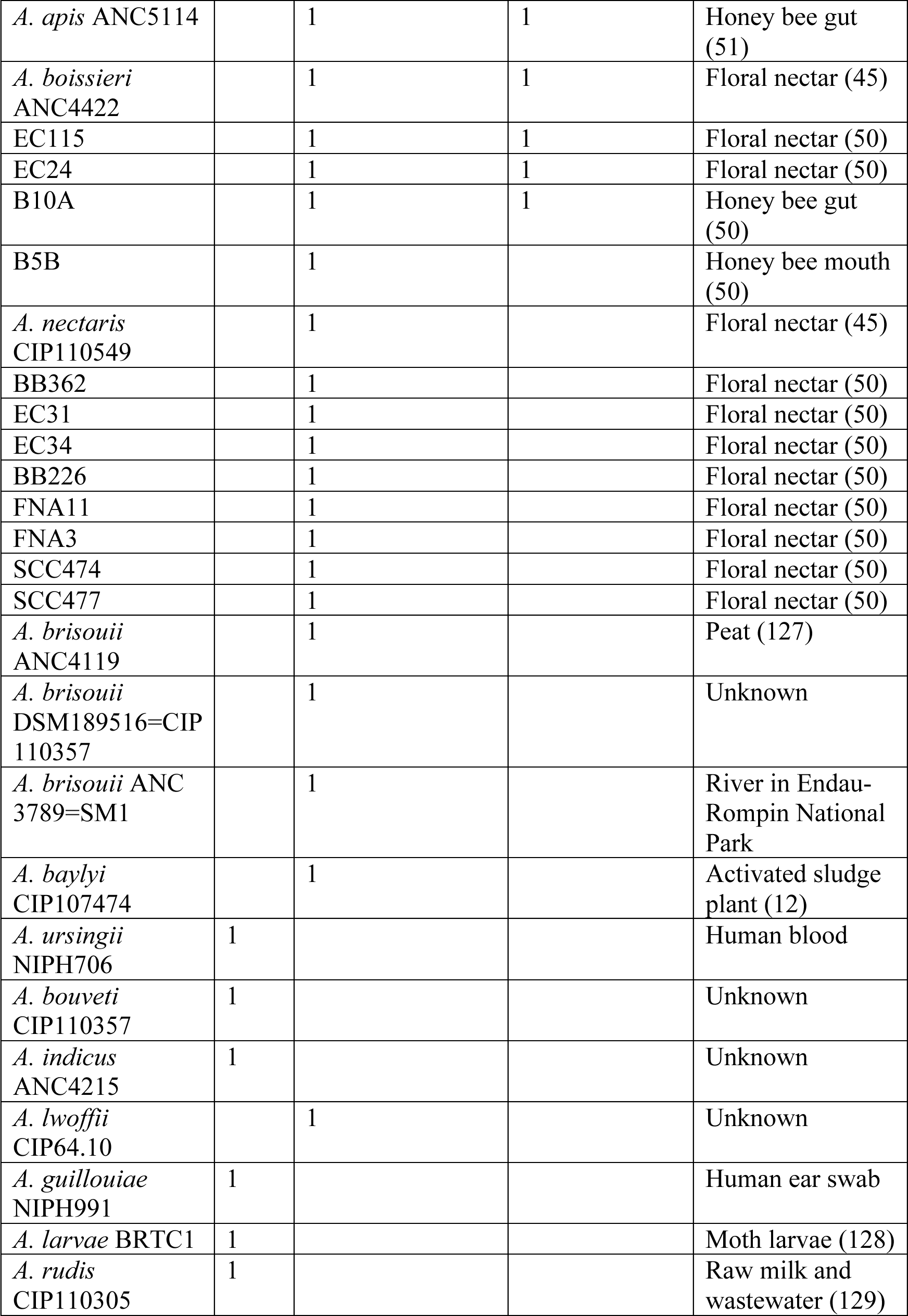
Presence and absence of phosphotransferase system (PTS) genes within *Acinetobacter* strains and the isolation environment for each strain (citations provided where available).

Because of the difference in sugars available to environmental versus nectar-dwelling *Acinetobacter*, we further investigated their carbohydrate metabolic capabilities by comparing carbohydrate active enzymes identified using the CAZY database (91, 92). Genomes of environmental species contain significantly more genes in the glycosyl transferase (GT) family (p-value = 0.001), with an average of 23 genes per environmental species genome and 15 per nectar dweller species genome (Fig. S3). This difference is mainly driven by enzymes in the GT2 and GT4 families. The role of these specific genes within *Acinetobacter* species is unclear, but generally enzymes in these families are involved in the synthesis of cell wall, capsular, and extracellular biofilm polysaccharides (93–96), suggesting that some of these functions may be different in nectar-dwelling *Acinetobacter*. We note, however, that several biofilm formation genes are found in the nectar-dwelling strains, including *pgaABCD* genes of the poly-β-1,6-*N*- acetyl-D-glucosamine (PGA) operon responsible for the maintenance of biofilm stability (97). Of the 15 nectar-dwelling strains, nine have the complete PGA operon, while four out of the 11 environmental strains have the full operon. Of the individual genes in the operon *A. pollinis* strains have 2-6 times the copies of *pgaB* genes, which contain binding domains critical for export of PGA (98), suggesting biofilm formation as an important trait in nectar-dwelling environments.

In contrast to the decrease in GT enzymes, nectar-dwellers contain significantly more genes in glycoside hydrolase (GH) families (p-value=8.152e-8), averaging 15 enzymes per species compared to approximately 13 genes per environmental species (Fig. S3). This pattern was mainly driven by genes in the GH 28 family, which are involved in the breakdown of the polygalacturonic acid backbone of pectin (99). Pectin is a major component of plant cells walls, and we hypothesize, as discussed below, that the ability to degrade this polysaccharide may be beneficial in floral nectar.

### Novel functions in nectar

We investigated which genes had been likely gained by HGT in nectar-dwelling *Acinetobacter*, with the hypothesis that such genes may confer novel functions. We found genomic islands within all members of the nectar-dwelling clade and some strains also contained intact prophages within their genomes. Gene count from genomic islands ranged from 111 – 352 genes with approximately 50% annotated as “hypothetical” and the remainder involved in plasmid or transposon mobilization, phage replication, or Type1 secretion components.

Mobilization genes were present in genomic islands within the nectar dwelling clade suggesting that movement of genomic material is facilitated by mobile elements like plasmids (Table S2).

Among known functions within genomic islands, we found genes coding for pectin degradation enzymes, specifically PL1 family pectin lyases and GH28 family polygalacturnonases. All species in the nectar-dwelling clade contain at least one of these genes, with several species possessing multiple copies of genes associated with the degradation of pectin (Table S3). Pectin is a recalcitrant polysaccharide that provides structural stability in plant cell walls and the outer layers of pollen grains (100). Among bacteria, enzymes for degrading pectin are commonly found in necrotrophic plant pathogens, which use them to digest plant tissue (101). These enzymes are notably absent among all *Acinetobacter* genomes in GenBank, with the exception of the orthologs found in the nectar clade (Table S3). Sequences in GenBank with the highest similarity to nectar-clade orthologs of PL1 and GH28 genes are found outside of the genus in plant pathogens such as *Pectobacterium*, *Erwinia* and *Dickeya* (Fig. 3A-B). This pattern supports that these genes were acquired by nectar-dwelling *Acinetobacter* by HGT, likely from a necrotrophic plant pathogen in the Enterobacterales.

**Figure 3.**
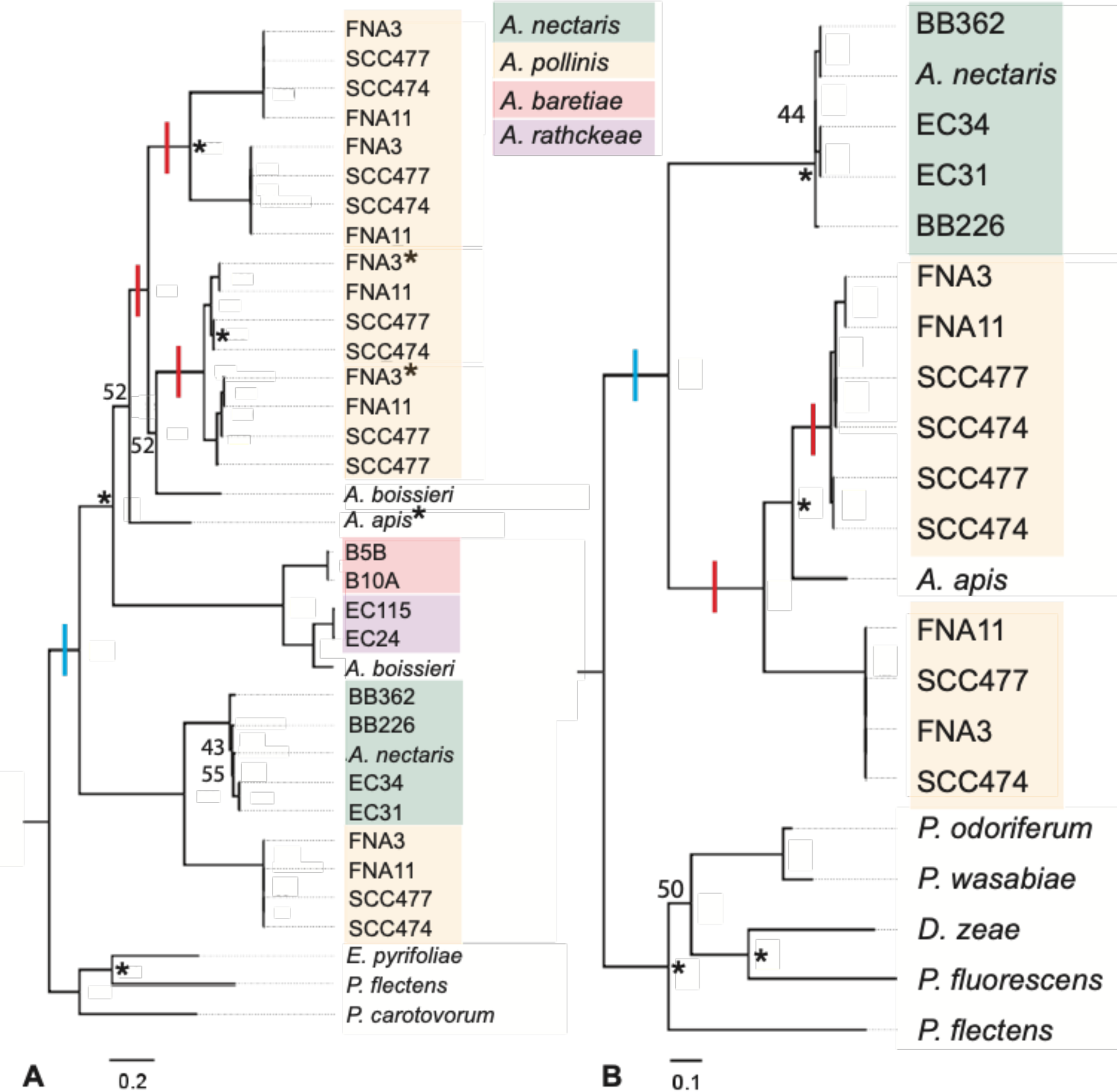
Maximum likelihood phylogenetic trees of A) the polygalacturonase gene, and B) the pectin lyase gene from nectar-dwelling *Acinetobacter* strains and the most closely related orthologs from plant pathogens. Red bars indicate likely duplication events and blue bars indicated horizontal gene transfer events. Bootstrap values below 80 are displayed at nodes. Asterisks show nodes and tips with significant positive selection. Orthologs from outside of *Acinetobacter* are from plant pathogens (*Erwinia pyrifoliae* (GenBank accession CP023567), *Pseudomonas flectans* (JAEE01000003), *Pectobacterium carotovorum* (CP001657), *Pectobacterium odoriferum* (MTAN01000008), *Pectobacterium wasabiae* (CP015750), *Dickeya zeae* (CP006929), and *Pseudomonas fluorescens* (OV986001)). Outgroup orthologs represent the best BLAST hits (>50% amino acid sequence identity, one ortholog per species) in GenBank databases.

Tracing the pattern of gains in pectin degradation genes onto the phylogeny of nectar-dwelling *Acinetobacter* suggests that at least one copy each of the pectin lyase and polygalacturonase genes were present in the common ancestor of the nectar-clade *Acinetobacter* species analyzed here (Fig. 3A-B). In several strains these two orthologs are located next to each other on the chromosome, so they may have been gained together in one event. As seen in the gene trees for the pectin lyase and polygalacturonase orthologs, we determined that these genes experienced multiple duplication events with paralogs sister to each other (Fig. 3A-B). These duplications occurred within the species *Acinetobacter pollinis*, which contains six copies of polygalacturonase and three copies of pectin lyase (within the strain SCC477 as an example). Additional horizontal transfers, losses, or duplication events may have occurred within the nectar clade, as some species have multiple copies within a gene tree (*A. boissieri*, Fig. 3A) or differences in topology between gene trees and species trees (*A. apis*, Fig. 3B). Some of these copies are on contigs that are likely from plasmids (based on increased read depth and the presence of plasmid replication genes), which may have facilitated duplication and transfers of these genes. Duplication was more common for the polygalacturonase than the pectin lyase genes, and the polygalacturonase genes were also the only example of multiple copies outside of the *A. pollinis* (Fig. 3A).

The fact that these genes have been maintained, and even duplicated, within the nectar-dwelling clade suggests that they may serve an important ecological role for these bacteria. In support of this, amino acid substitutions in several of pectin degrading enzyme protein sequences show signatures of positive selection (Table S4). Positive selection was detected at the nodes and tips of the polygalacturonase gene tree (Fig. 3), particularly for *A. pollinis* (nine sites) and *A. apis* (seven sites) orthologs (Table S4). The high number of duplication events of these genes in *A. pollinis*, together with signatures of positive selection, suggests that pectin degrading enzymes may be functionally diversifying in this species. Both of these enzymes cleave linkages in the polygalacturonic acid backbone of pectin (102). Necrotrophic plant pathogens typically have diverse copies of these enzymes, with slight variations in catalytic ability, in order to effectively degrade pectin (102–104). Potentially, this pattern is convergently evolving in *A. pollinis*.

To investigate the potential function of the amino acids under selection in *Acinetobacter* polygalacturonase enzymes, we generated predicted protein structures of representative orthologs from *A. pollinis* strain FNA3 and the *A. apis* type strain (105). The polygalacturonase binding site contains four highly conserved motifs NTD, G/QDD, G/SHG and RIK. NTD and RIK are highly conserved catalytic regions while G/QDD and G/SHG are substrate binding regions (106). In nectar strain orthologs all conserved regions were present (NTD, GDD, GHG, RIK) but no sites under selection were near these active sites (Fig. 4). It is therefore unclear from their location on the protein structure how sites under selection might impact protein function.

**Figure 4:**
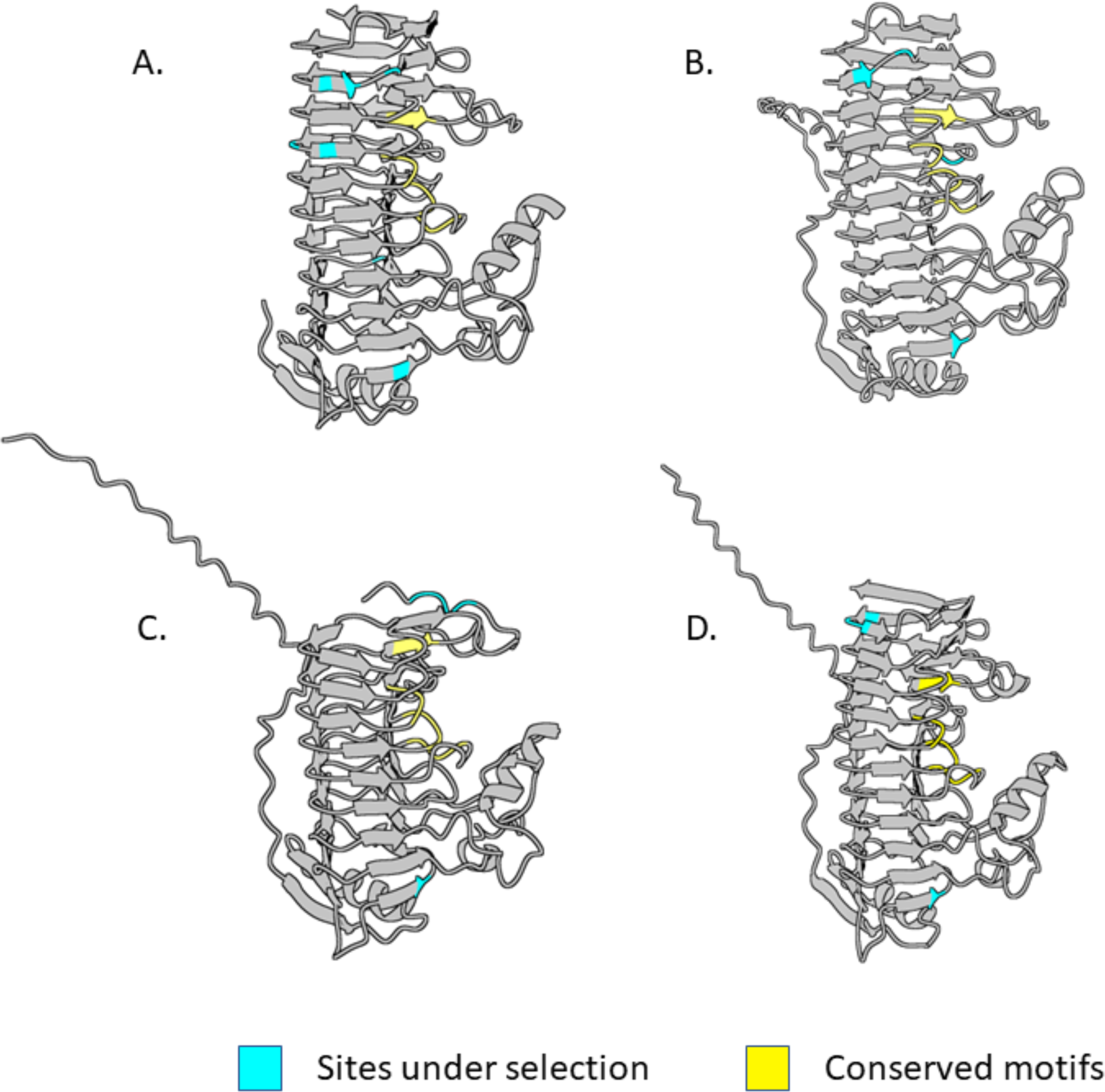
Alphafold protein predictions of polygalacturonase genes from *Acinetobacter apis* (A)*, Acinetobacter pollinis* FNA3 (B-C), and *Phaseolibacter flectens* (D). Conserved motifs and sites under selection are highlighted blue and yellow respectively. Secretion signal tags are present in B-D. These extend off the protein to the left in C-D, but backwards in B. B is rotated in Fig. S2 to show the secretion signal.

It is not known how much nectar-dwelling *Acinetobacter* interact with major sources of pectin in plant tissue, but to our knowledge they have not been observed to infect plants.

However, microbes in nectar are regularly interact with pollen grains, which are introduced into nectar by pollinator activity (107). In fact, some *Acinetobacter* strains can cause pollen grains to burst open or pseudogerminate (29). This ability is beneficial for *Acinetobacter*, as it is associated with increased growth in nectar when pollen is present (29). Floral nectar has been shown to be nitrogen limiting for both yeasts and bacteria (80, 81), so the ability to access nitrogen from pollen within nectar could increase microbial fitness. However, pollen is protected by a resistant exine layer and is difficult to degrade (108). Pectin is an essential component of pollen cell walls and pollen tubes (109), and pectin degrading enzymes have been hypothesized to be involved in pollen breakdown by bacterial gut symbionts of honey bees (110). We hypothesize that the pectin degradation enzymes in *Acinetobacter* could be involved in accessing nutrients from pollen, which could explain the apparent importance of genes coding for such enzymes in the clade. In support of this, we find that most of the GH28 and PL1 proteins in *Acinetobacter* have secretion signals (Fig. 4B-D, Fig. S4B-D), similar to secreted pectin degrading enzymes in *Pectobacterium* (102), suggesting that they should act extracellularly. Furthermore, we find the most selection on these genes within *A. pollinis* and *A. apis*. The former shows strong impacts on pollen bursting and pseudogermination (29) and the latter was isolated from honey bees and is likely to encounter pollen regularly. We speculate that the ability to degrade pectin could be a key trait allowing *Acinetobacter* to thrive in nectar and in association with pollinators.

### Conclusions

In this study we show that the ecological change from soil-dwelling to nectar-dwelling led to genomic reduction, possibly due to genome streamlining, a phenomenon common among bacteria living in oligotrophic habitats (66). Even with this streamlining, nectar-dwelling species had higher levels of gene gain events compared to gene losses. Many of the gain gains occur at early branching nodes within the nectar clade, and the nectar clade had an increase in the number of genes involved in monosacharride metabolism and sugar transport, likely due to the high sugar environment of nectar (78). We also found changes in nitrogen and amino acids metabolism genes suggesting a switch towards nitrogen scavenging in comparison to environmental *Acinetobacter*, consistent with nitrogen limitation in floral nectar (55). Nectar-dwelling *Acinetobacter* species have also acquired pectin degrading enzymes, presumably through HGT, from plant pathogens. We found duplication, diversification, and positive selection within pectin degrading genes, supporting our hypothesis that these genes may provide an important ecological function. Overall, we find that genome evolution from gene loss, diversification, and HGT may have all contributed to the *Acinetobacter* habitat switch to floral nectar.

## Materials and Methods

### Phylogenetic analyses

A phylogenomic species tree was inferred using 26 *Acinetobacter* genomes and *Pseudomonas syringae* pv. *tomato* strain DC3000 as an outgroup (Table 1). These included sequences from nine environmental *Acinetobacter* species, and three previously published nectar-dwelling and pollinator associated *Acinetobacter* genomes (45, 51) obtained from GenBank. We did not include strains from the clades containing the animal pathogens *A. baumannii* or *A. parvus*, as previous work found these groups to have undergone distinct evolutionary changes compared to soil-dwelling relatives (4). Our recently published nectar-dwelling *Acinetobacter* genomes (see (50) for isolation information) were assembled using Discovar *de novo* (111), checked for completeness using CheckM (112), and annotated using the RAST Server (112) Nucleotide sequences from previously published *Acinetobacter* genomes were downloaded from GenBank and annotated in RAST for consistency. Protein sequences were used in the PhyloPhlAn 3.0 pipeline (114) to determine conserved proteins within *Acinetobacter* genomes. PhyloPhlAn identified 399 conserved proteins and their nucleotide sequences were used for phylogenetic tree reconstruction. Nucleotide sequences for the identified protein sequences were aligned using MAFFT (115). Maximum likelihood trees were reconstructed using IQ-Tree (116) with bootstrapping set to 1000 and a symmetric substitution model (117). Welch’s t-test was used to determine the significance between genome size from the nectar clade and environmental clade.

Gene trees were reconstructed using polygalacturonase and pectin lyase genes from the *Acinetobacter* genomes. Outgroups were selected by using nectar-dwelling *Acinetobacter* spp. pectin lyase and polygalacturonase genes as BLAST queries in GenBank. We found that all of the best BLAST hits for these genes were from necrotrophic plant pathogens so we included the most similar representative sequences in the analyses. *Acinetobacter* polygalacturonase and pectin lyase genes were identified in our newly sequenced genomes by RAST annotation, BLAST of the genomes using plant pathogen orthologs as queries, and comparison with the CAZy database to confirm that we had identified all orthologs. Genes were aligned using MAFFT (115) and were used for maximum likelihood phylogenetic inference in IQ-Tree using the TIM3 substitution model which was selected using model finder in IQ-Tree.

### Ortholog analyses

Orthologous protein sequence clustering was conducted using OrthoMCL (75) using the following parameters: mode = 1, inflation = 2, pi cutoff = 50. To determine the ancestral state of orthologs across the *Acinetobacter* species tree, the software package Count was used (118) implementing Wagner Parsimony. This analysis determines the genes gained and lost at each node of the *Acinetobacter* phylogenomic tree minimizing state changes and assuming all character states are reversible (119). To estimate the number of pseudogenes present within the genomes, we used the program Pseudofinder (120). The algorithm identifies pseudogenes from Genbank files by analyzing average coding sequence (CDS) length, fragmented CDS, and intergenic pseudogenes and alignment lengths are compared against homologs identified by blastp hits from a reference protein database (121). The following parameters were used to predict potential intergenic, fragmented, truncated, and long pseudogenes: intergenic length = 30, length pseudo = 0.65, shared hits = 0.5, hitcap = 15, intergenic threshold = 0.3. Plasmids were assembled using unfiltered paired end 2 x 250 Illumina sequencing reads (50) and the plasmidSPAdes assembly tool (122). Plasmids less than 1000 bp were excluded from final plasmid count. Fisher’s exact test was used to calculate all significance test between orthologs from the nectar groups and the environmental group.

### Selection and protein analyses

dN/dS (ω) values were estimated for the polygalacturonase and pectin lyase coding region alignments and IQ gene tree phylogenies using codeml in the PAML v4.4 package, with gaps included (123). For each branch and node in the phylogenies, a likelihood-ratio test for positive selection was performed to compare nested branch-site models (Model Anull versus Model A) (124) (Table S4).

The AI protein prediction software, Alphafold, was used to predict protein structure of pectin degradation enzymes. The Alphafold algorithm is a neural network that generates a multiple sequence alignment (MSA) from the query protein sequence provided and extracts evolutionary information to generate protein predictions (105). A web version of Alphafold was used for these predictions (125). In order to determine if amino acid sites under selection were functionally important, we predicted the structure of polygalacturonase genes identified to be under positive selection from the PAML analysis, specifically genes from *Acinetobacter pollinis* strain FNA3 (GenBank locus tags I2F29_RS12745 and I2F29_RS12925), *Acinetobacter apis* (CFY84_RS01715), and *Phaseolibacter flectens* (L871_RS0110380). Three-dimensional protein predictions were edited using ChimeraX to highlight sites under selection (126).

## Data availability

All data is available in the supplemental materials or NCBI databases. Genomic assemblies are available in GenBank under accessions VTDP00000000, VTDO00000000, VTDM00000000, VTDL00000000, JAEQDL000000000, JAERJC000000000, JAERJB000000000, JAEQDM000000000, VTDS00000000, VTDT00000000, VTDR00000000, and VTDQ00000000. Sequence reads are available in the SRA under accessions SRR26518995- SRR26519002.

## Acknowledgements

We thank the Cornell Biotechnology Resource Center for sequencing and Heather Feaga for consultation on analyses. We are grateful to Rachel Vannette, Tadashi Fukami, and Bart Lievens for sharing isolates for sequencing.

Support was provided by the SUNY Graduate Diversity Fellowship (V.A.S.) and the Cornell Atkinson Center for Sustainability Sustainable Biodiversity Fund (V.A.S.).

V.A.S. and T.A.H. conceived of the study and V.A.S., T.A.H., L.J.B., and T.R. performed analyses. Writing was collaborative between V.A.S., T.A.H., and T.R., with comments from L.J.B.

## References

1. Tan SYY, Chua SL, Liu Y, Høiby N, Andersen LP, Givskov M, Song Z, Yang L. 2013. Comparative Genomic Analysis of Rapid Evolution of an Extreme-Drug-Resistant Acinetobacter baumannii Clone. Genome Biol Evol 5:807–818.

2. Zarrilli R, Pournaras S, Giannouli M, Tsakris A. 2013. Global evolution of multidrug-resistant Acinetobacter baumannii clonal lineages. Int J Antimicrob Agents 41:11–19.

3. Adams MD, Chan ER, Molyneaux ND, Bonomo RA. 2010. Genomewide analysis of divergence of antibiotic resistance determinants in closely related isolates of Acinetobacter baumannii. Antimicrob Agents Chemother 54:3569–3577.

4. Garcia-Garcera M, Touchon M, Brisse S, Rocha EPC. 2017. Metagenomic assessment of the interplay between the environment and the genetic diversification of *Acinetobacter*. Environ Microbiol 19:5010–5024.

5. Harding CM, Hennon SW, Feldman MF. 2018. Uncovering the mechanisms of Acinetobacter baumannii virulence. Nat Rev Microbiol 16:91.

6. Imperi F, Antunes LCS, Blom J, Villa L, Iacono M, Visca P, Carattoli A. 2011. The genomics of Acinetobacter baumannii: Insights into genome plasticity, antimicrobial resistance and pathogenicity. IUBMB Life 63:1068–1074.

7. Koskella B, Vos M. 2015. Adaptation in Natural Microbial Populations. Annu Rev Ecol Evol Syst 46:503–522.

8. Touchon M, Cury J, Yoon EJ, Krizova L, Cerqueira GC, Murphy C, Feldgarden M, Wortman J, Clermont D, Lambert T, Grillot-Courvalin C, Nemec A, Courvalin P, Rocha EPC. 2014. The genomic diversification of the whole Acinetobacter genus: Origins, mechanisms, and consequences. Genome Biol Evol 6:2866–2882.

9. Park J, Park W. 2011. Phenotypic and physiological changes in Acinetobacter sp. strain DR1 with exogenous plasmid. Curr Microbiol 62:249–254.

10. Traglia GM, Chua K, Centron D, Tolmasky ME, Ramírez MS. 2014. Whole-Genome Sequence Analysis of the Naturally Competent Acinetobacter baumannii Clinical Isolate A118. Genome Biol Evol 6:2235–2239.

11. Oliveira PH, Touchon M, Cury J, Rocha EPC. 2017. The chromosomal organization of horizontal gene transfer in bacteria. Nature Communications 2017 8:1 8:1–11.

12. Santala S, Santala V. 2021. Acinetobacter baylyi ADP1-naturally competent for synthetic biology. Essays Biochem 65:309–318.

13. Veress A, Nagy T, Wilk T, Kömüves J, Olasz F, Kiss J. 2020. Abundance of mobile genetic elements in an Acinetobacter lwoffii strain isolated from Transylvanian honey sample. Scientific Reports 2020 10:1 10:1–14.

14. Fondi M, Bacci G, Brilli M, Papaleo MC, Mengoni A, Vaneechoutte M, Dijkshoorn L, Fani R. 2010. Exploring the evolutionary dynamics of plasmids: The Acinetobacter pan- plasmidome. BMC Evol Biol 10:1–15.

15. Blaschke U, Skiebe E, Kaatz M, Higgins PG, Pfeifer Y, Wilharm G. 2018. Complete Genome Sequencing of Acinetobacter sp. Strain LoGeW2-3, Isolated from the Pellet of a White Stork, Reveals a Novel Class D Beta-Lactamase Gene. Genome Announc 6.

16. Kim DH, Jung SI, Kwon KT, Ko KS. 2017. Occurrence of diverse AbGRI1-type genomic islands in Acinetobacter baumannii global clone 2 isolates from South Korea. Antimicrob Agents Chemother 61.

17. Sezmis AL, Woods LC, Peleg AY, McDonald MJ. 2023. Horizontal Gene Transfer, Fitness Costs and Mobility Shape the Spread of Antibiotic Resistance Genes into Experimental Populations of Acinetobacter Baylyi. Mol Biol Evol 40.

18. Salto IP, Torres Tejerizo G, Wibberg D, Pühler A, Schlüter A, Pistorio M. 2018. Comparative genomic analysis of Acinetobacter spp. plasmids originating from clinical settings and environmental habitats. Sci Rep 8:7783.

19. Treangen TJ, Rocha EPC. 2011. Horizontal Transfer, Not Duplication, Drives the Expansion of Protein Families in Prokaryotes. PLoS Genet 7:e1001284.

20. Norton MD, Spilkia AJ, Godoy VG. 2013. Antibiotic resistance acquired through a DNA damage-inducible response in Acinetobacter baumannii. J Bacteriol 195:1335–1345.

21. Kassen R. 2019. Experimental Evolution of Innovation and Novelty. Trends Ecol Evol 34:712–722.

22. Kondrashov FA. 2012. Gene duplication as a mechanism of genomic adaptation to a changing environment. Proceedings of the Royal Society B: Biological Sciences. Proc R Soc B 279:1108.

23. Bolhuis H, Severin I, Confurius-Guns V, Wollenzien UIA, Stal LJ. 2009. Horizontal transfer of the nitrogen fixation gene cluster in the cyanobacterium Microcoleus chthonoplastes. The ISME Journal 2010 4:1 4:121–130.

24. Colombi E, Hill Y, Lines R, Sullivan JT, Kohlmeier MG, Christophersen CT, Ronson CW, Terpolilli JJ, Ramsay JP. 2023. Population genomics of Australian indigenous Mesorhizobium reveals diverse nonsymbiotic genospecies capable of nitrogen-fixing symbioses following horizontal gene transfer. Microb Genom 9:000918.

25. Pacheco S, Gómez I, Chiñas M, Sánchez J, Soberón M, Bravo A. 2021. Whole Genome Sequencing Analysis of Bacillus thuringiensis GR007 Reveals Multiple Pesticidal Protein Genes. Front Microbiol 12:3268.

26. Fournier P-E, Vallenet D, Barbe V, Audic S, Ogata H, Poirel L, Richet H, Robert C, Mangenot S, Abergel C, Nordmann P, Weissenbach J, Raoult D, Claverie J-M. 2006. Comparative Genomics of Multidrug Resistance in Acinetobacter baumannii. PLoS Genet 2:e7.

27. Savory EA, Fuller SL, Weisberg AJ, Thomas WJ, Gordon MI, Stevens DM, Creason AL, Belcher MS, Serdani M, Wiseman MS, Grünwald NJ, Putnam ML, Chang JH. 2017. Evolutionary transitions between beneficial and phytopathogenic rhodococcus challenge disease management. Elife 6.

28. Melnyk RA, Hossain SS, Haney CH. 2019. Convergent gain and loss of genomic islands drive lifestyle changes in plant-associated Pseudomonas. The ISME Journal 2019 13:6 13:1575–1588.

29. Christensen SM, Munkres I, Vannette RL. 2021. Nectar bacteria stimulate pollen germination and bursting to enhance their fitness Short Title: Nectar bacteria digest pollen via induced germination. Current Biology 31:4373–4380.

30. Nicolson SW, Fleming PA. 2003. Nectar as food for birds: the physiological consequences of drinking dilute sugar solutions. Plant Systematics and Evolution 238:139–153.

31. Pamminger T, Becker R, Himmelreich S, Schneider CW, Bergtold M. 2019. The nectar report: Quantitative review of nectar sugar concentrations offered by bee visited flowers in agricultural and non-agricultural landscapes. PeerJ 2019:e6329.

32. Martin VN, Schaeffer RN, Fukami T. 2022. Potential effects of nectar microbes on pollinator health. Philosophical Transactions of the Royal Society B 377.

33. Russell KA, McFrederick QS. 2022. Elevated Temperature May Affect Nectar Microbes, Nectar Sugars, and Bumble Bee Foraging Preference. Microb Ecol 84:473–482.

34. Pozo MI, Jacquemyn H. 2019. Addition of pollen increases growth of nectar-living yeasts. FEMS Microbiol Lett 366:191.

35. Pozo MI, Lachance MA, Herrera CM. 2012. Nectar yeasts of two southern Spanish plants: the roles of immigration and physiological traits in community assembly. FEMS Microbiol Ecol 80:281–293.

36. Lievens B, Hallsworth JE, Pozo MI, Belgacem Z ben, Stevenson A, Willems KA, Jacquemyn H. 2015. Microbiology of sugar-rich environments: Diversity, ecology and system constraints. Environ Microbiol 17:278–298.

37. Herrera CM, Canto A, Pozo MI, Bazaga P. 2010. Inhospitable sweetness: nectar filtering of pollinator-borne inocula leads to impoverished, phylogenetically clustered yeast communities. Proceedings of the Royal Society B: Biological Sciences 277:747–754.

38. Herrera CM. 2017. Scavengers that fit beneath a microscope lens. Ecology 98:2725–2726.

39. Levin E, McCue MD, Davidowitz G. 2017. More than just sugar: allocation of nectar amino acids and fatty acids in a Lepidopteran. Proceedings of the Royal Society B: Biological Sciences 284.

40. Dhami MK, Hartwig T, Fukami T. 2016. Genetic basis of priority effects: insights from nectar yeast. Proceedings of the Royal Society B: Biological Sciences 283.

41. Herrera CM. 2017. Scavengers that fit beneath a microscope lens. Ecology 98:2725–2726.

42. Schaeffer RN, Vannette RL, Brittain C, Williams NM, Fukami T. 2017. Non-target effects of fungicides on nectar-inhabiting fungi of almond flowers. Environ Microbiol Rep 9:79– 84.

43. Fridman S, Izhaki I, Gerchman Y, Halpern M. 2012. Bacterial communities in floral nectar. Environ Microbiol Rep 4:97–104.

44. Morris MM, Frixione NJ, Burkert AC, Dinsdale EA, Vannette RL. 2020. Microbial abundance, composition, and function in nectar are shaped by flower visitor identity. FEMS Microbiol Ecol 96:fiaa003.

45. Álvarez-Pérez S, Lievens B, Jacquemyn H, Herrera CM. 2013. Acinetobacter nectaris sp. nov. and Acinetobacter boissieri sp. nov., isolated from floral nectar of wild Mediterranean insect-pollinated plants. Int J Syst Evol Microbiol 63:1532–1539.

46. Álvarez-Pérez S, de Vega C, Pozo MI, Lenaerts M, Van Assche A, Herrera CM, Jacquemyn H, Lievens B. 2016. Nectar yeasts of the Metschnikowia clade are highly susceptible to azole antifungals widely used in medicine and agriculture. FEMS Yeast Res 16.

47. Aizenberg-Gershtein Y, Izhaki I, Halpern M. 2013. Do Honeybees Shape the Bacterial Community Composition in Floral Nectar? PLoS One 8:e67556.

48. Samuni-Blank M, Izhaki I, Laviad S, Bar-Massada A, Gerchman Y, Halpern M. 2014. The Role of Abiotic Environmental Conditions and Herbivory in Shaping Bacterial Community Composition in Floral Nectar. PLoS One 9:e99107.

49. von Arx M, Moore A, Davidowitz G, Arnold AE. 2019. Diversity and distribution of microbial communities in floral nectar of two night-blooming plants of the Sonoran Desert. PLoS One 14:e0225309.

50. Alvarez-Perez S, Baker LJ, Morris MM, Tsuji K, Sanchez VA, Fukami T, Vannette RL, Lievens B, Hendry TA. 2021. Acinetobacter pollinis sp. nov., Acinetobacter baretiae sp. nov. and Acinetobacter rathckeae sp. nov., isolated from floral nectar and honey bees. Int J Syst Evol Microbiol 71:004783.

51. Kim PS, Shin N-R, Kim JY, Yun J-H, Hyun D-W, Bae J-W. 2014. Acinetobacter apis sp. nov., isolated from the intestinal tract of a honey bee, Apis mellifera. Journal of Microbiology 52:639–645.

52. McFrederick QS, Rehan SM, MC FREDERICK QS, Rehan SM. 2016. Characterization of pollen and bacterial community composition in brood provisions of a small carpenter bee. Mol Ecol 25:2302–2311.

53. Tang QH, Miao CH, Chen YF, Dong ZX, Cao Z, Liao SQ, Wang JX, Wang ZW, Guo J. 2021. The composition of bacteria in gut and beebread of stingless bees (Apidae: Meliponini) from tropics Yunnan, China. Antonie van Leeuwenhoek, International Journal of General and Molecular Microbiology 114:1293–1305.

54. Evans JD, Armstrong TN. 2015. Inhibition of the American foulbrood bacterium, Paenibacillus larvae larvae, by bacteria isolated from honey bees. J Apiculture Research 44: 168–171.

55. Morales-Poole JR, de Vega C, Tsuji K, Jacquemyn H, Junker RR, Herrera CM, Michiels C, Lievens B, Álvarez-Pérez S. 2022. Sugar Concentration, Nitrogen Availability, and Phylogenetic Factors Determine the Ability of Acinetobacter spp. and Rosenbergiella spp. to Grow in Floral Nectar. Microb Ecol 1:1–15.

56. Sharaby Y, Rodríguez-Martínez S, Lalzar M, Halpern M, Izhaki I. 2020. Geographic partitioning or environmental selection: What governs the global distribution of bacterial communities inhabiting floral nectar? Science of The Total Environment 749:142305.

57. Vannette RL, Fukami T. 2017. Dispersal enhances beta diversity in nectar microbes. Ecol Lett 20:901–910.

58. Lee C, Tell LA, Hilfer T, Vannette RL. 2019. Microbial communities in hummingbird feeders are distinct from floral nectar and influenced by bird visitation. Proceedings of the Royal Society B: Biological Sciences 286:20182295.

59. Simonsen AK. 2021. Environmental stress leads to genome streamlining in a widely distributed species of soil bacteria. The ISME Journal 2021 16:2 16:423–434.

60. Giovannoni SJ, Tripp HJ, Givan S, Podar M, Vergin KL, Baptista D, Bibbs L, Eads J, Richardson TH, Noordewier M, Rappé MS, Short JM, Carrington JC, Mathur EJ. 2005. Genetics: Genome streamlining in a cosmopolitan oceanic bacterium. Science (1979) 309:1242–1245.

61. Giovannoni SJ, Cameron Thrash J, Temperton B. 2014. Implications of streamlining theory for microbial ecology. The ISME Journal 2014 8:8 8:1553–1565.

62. Liang J, Liu J, Wang X, Sun H, Zhang Y, Ju F, Thompson F, Zhang XH. 2022. Genomic Analysis Reveals Adaptation of Vibrio campbellii to the Hadal Ocean. Appl Environ Microbiol 88.

63. Lee M-C, Marx CJ. 2012. Repeated, Selection-Driven Genome Reduction of Accessory Genes in Experimental Populations. PLoS Genet 8:e1002651.

64. Karcagi I, Draskovits G, Umenhoffer K, Fekete G, Kovács K, Méhi O, Balikó G, Szappanos B, Györfy Z, Fehér T, Bogos B, Blattner FR, Pál C, Pósfai G, Papp B. 2016. Indispensability of Horizontally Transferred Genes and Its Impact on Bacterial Genome Streamlining. Mol Biol Evol 33:1257–1269.

65. Martinez-Gutierrez CA, Aylward FO, Lerat E. 2019. Strong Purifying Selection Is Associated with Genome Streamlining in Epipelagic Marinimicrobia. Genome Biol Evol 11:2887–2894.

66. Simonsen AK. 2022. Environmental stress leads to genome streamlining in a widely distributed species of soil bacteria. ISME J 16:423–434.

67. Sabath N, Ferrada E, Barve A, Wagner A. 2013. Growth Temperature and Genome Size in Bacteria Are Negatively Correlated, Suggesting Genomic Streamlining During Thermal Adaptation. Genome Biol Evol 5:966–977.

68. Moran NA. 1996. Accelerated evolution and Muller’s rachet in endosymbiotic bacteria. Proceedings of the National Academy of Sciences 93:2873–2878.

69. Hendry T, Freed L, Dana F, Danté F, Sutton T, Lopez J. 2018. Ongoing Transposon- Mediated Genome Reduction in the Luminous Bacterial Symbionts of Deep-Sea Ceratioid Anglerfishes. mBio 9:e01033–18.

70. Hendry T, de Wet J, Dougan K, Dunlap P. 2016. Genome Evolution in the Obligate but Environmentally Active Luminous Symbionts of Flashlight Fish. Genome Biol Evol 8:2203–2213.

71. McCutcheon JP, Moran NA. 2012. Extreme genome reduction in symbiotic bacteria. Nat Rev Microbiol 10:13–26.

72. Angst P, Ameline C, Haag CR, Ben-Ami F, Ebert D, Fields PD. 2022. Genetic Drift Shapes the Evolution of a Highly Dynamic Metapopulation. Mol Biol Evol 39.

73. Repizo GD, Espariz M, Seravalle JL, Miloslavich JID, Steimbrüch BA, Shuman HA, Viale AM. 2020. Acinetobacter baumannii NCIMB8209: a Rare Environmental Strain Displaying Extensive Insertion Sequence-Mediated Genome Remodeling Resulting in the Loss of Exposed Cell Structures and Defensive Mechanisms. mSphere 5.

74. Vallenet D, Nordmann P, Barbe V, Poirel L, Mangenot S, Bataille E, Dossat C, Gas S, Kreimeyer A, Lenoble P, Oztas S, Poulain J, Segurens B, Robert C, Abergel C, Claverie JM, Raoult D, Médigue C, Weissenbach J, Cruveiller S. 2008. Comparative Analysis of Acinetobacters: Three Genomes for Three Lifestyles. PLoS One 3:e1805.

75. Li L, Stoeckert CJ, Roos DS. 2003. OrthoMCL: Identification of Ortholog Groups for Eukaryotic Genomes. Genome Res 13:2178.

76. Zhou Z, Tran PQ, Kieft K, Anantharaman K. 2020. Genome diversification in globally distributed novel marine Proteobacteria is linked to environmental adaptation. The ISME Journal 2020 14:8 14:2060–2077.

77. Kolosova N, Gorenstein N, Kish CM, Dudareva N. 2001. Regulation of Circadian Methyl Benzoate Emission in Diurnally and Nocturnally Emitting Plants. Plant Cell 13:2333– 2347.

78. Wolff D. 2006. Nectar sugar composition and volumes of 47 species of Gentianales from a southern Ecuadorian montane forest. Ann Bot 97.

79. Pozo MI, Herrera CM, Van den Ende W, Verstrepen K, Lievens B, Jacquemyn H. 2015. The impact of nectar chemical features on phenotypic variation in two related nectar yeasts. FEMS Microbiol Ecol 91.

80. Morales-Poole JR, de Vega C, Tsuji K, Jacquemyn H, Junker RR, Herrera CM, Michiels C, Lievens B, Álvarez-Pérez S. 2022. Sugar Concentration, Nitrogen Availability, and Phylogenetic Factors Determine the Ability of Acinetobacter spp. and Rosenbergiella spp. to Grow in Floral Nectar. Microb Ecol 86:377–391.

81. Álvarez-Pérez S, Tsuji K, Donald M, Van Assche A, Vannette RL, Herrera CM, Jacquemyn H, Fukami T, Lievens B. 2021. Nitrogen Assimilation Varies Among Clades of Nectar- and Insect-Associated Acinetobacters. Microb Ecol 81:990–1003.

82. Chalcoff VR, Aizen MA, Galetto L. 2006. Nectar Concentration and Composition of 26 Species from the Temperate Forest of South America. Ann Bot 97:413–421.

83. Arvay E, Biggs BW, Guerrero L, Jiang V, Tyo K. 2021. Engineering Acinetobacter baylyi ADP1 for mevalonate production from lignin-derived aromatic compounds. Metab Eng Commun 13:e00173.

84. Durot M, le Fèvre F, de Berardinis V, Kreimeyer A, Vallenet D, Combe C, Smidtas S, Salanoubat M, Weissenbach J, Schachter V. 2008. Iterative reconstruction of a global metabolic model of Acinetobacter baylyi ADP1 using high-throughput growth phenotype and gene essentiality data. BMC Syst Biol 2:1–23.

85. Salcedo-Vite K, Sigala JC, Segura D, Gosset G, Martinez A. 2019. Acinetobacter baylyi ADP1 growth performance and lipid accumulation on different carbon sources. Appl Microbiol Biotechnol 103:6217–6229.

86. Orchard LMD, Kornberg HL. 1997. Sequence similarities between the gene specifying 1- phosphofructokinase (fruK), genes specifying other kinases in Escherichia coli K12, and lacC of Staphylococcus aureus. Proc R Soc Lond B Biol Sci 242:87–90.

87. Collins T, Gerday C, Feller G. 2005. Xylanases, xylanase families and extremophilic xylanases. FEMS Microbiol Rev 29:3–23.

88. Lengeler JW, Jahreis K. 2009. Bacterial PEP-dependent carbohydrate: phosphotransferase systems couple sensing and global control mechanisms. Contrib Microbiol 16:65–87.

89. Saier Jr. MH. 2015. The Bacterial Phosphotransferase System: New Frontiers 50 Years after Its Discovery. Microb Physiol 25:73–78.

90. Saier Jr. MH. 2015. The Bacterial Phosphotransferase System: New Frontiers 50 Years after Its Discovery. Microb Physiol 25:73–78.

91. Yin Y, Mao X, Yang J, Chen X, Mao F, Xu Y. 2012. dbCAN: a web resource for automated carbohydrate-active enzyme annotation. Nucleic Acids Res 445–451.

92. Drula E, Garron M-L, Dogan S, Lombard V, Henrissat B, Terrapon N. 2022. The carbohydrate-active enzyme database: functions and literature. Nucleic Acids Res 50:D571–D577.

93. Coutinho PM, Deleury E, Davies GJ, Henrissat B. 2003. An evolving hierarchical family classification for glycosyltransferases. J Mol Biol 328:307–317.

94. Akoolo L, Pires S, Kim J, Parker D. 2022. The Capsule of Acinetobacter baumannii Protects against the Innate Immune Response. J Innate Immun 14:543–554.

95. Harding CM, Hennon SW, Feldman MF. 2018. Uncovering the mechanisms of Acinetobacter baumannii virulence. Nat Rev Microbiol 16:91–102.

96. Zhang G, Baidin V, Pahil KS, Moison E, Tomasek D, Ramadoss NS, Chatterjee AK, McNamara CW, Young TS, Schultz PG, Meredith TC, Kahne D. 2018. Cell-based screen for discovering lipopolysaccharide biogenesis inhibitors. Proceedings of the National Academy of Sciences 115:6834–6839.

97. Yoshikane I, D RJ, Carlos G, Archana P, Jeannette T, Jeffrey M, J BT, F PJ, Tony R. 2008. Roles of pgaABCD Genes in Synthesis, Modification, and Export of the Escherichia coli Biofilm Adhesin Poly-β-1,6-N-Acetyl-d-Glucosamine. J Bacteriol 190:3670–3680.

98. Little DJ, Pfoh R, Le Mauff F, Bamford NC, Notte C, Baker P, Guragain M, Robinson H, Pier GB, Nitz M, Deora R, Sheppard DC, Howell PL. 2018. PgaB orthologues contain a glycoside hydrolase domain that cleaves deacetylated poly-β(1,6)-N-acetylglucosamine and can disrupt bacterial biofilms. PLoS Pathog 14.

99. Henrissat B, Davies G. 1997. Structural and sequence-based classification of glycoside hydrolases. Curr Opin Struct Biol 7:637–644.

100. Cankar K, Kortstee A, Toonen MAJ, Wolters-Arts M, Houbein R, Mariani C, Ulvskov P, Jorgensen B, Schols HA, Visser RGF, Trindade LM. 2014. Pectic arabinan side chains are essential for pollen cell wall integrity during pollen development. Plant Biotechnology Journal 12:492–502.

101. Jayani RS, Shukla SK, Gupta R. 2010. Screening of Bacterial Strains for Polygalacturonase Activity: Its Production by Bacillus sphaericus (MTCC 7542). Enzyme Res 2010.

102. Abbott DW, Boraston AB. 2008. Structural Biology of Pectin Degradation by Enterobacteriaceae. Microbiol Mol Biol Rev 72:301.

103. Sprockett DD, Piontkivska H, Blackwood CB. 2011. Evolutionary analysis of glycosyl hydrolase family 28 (GH28) suggests lineage-specific expansions in necrotrophic fungal pathogens. Gene 479:29–36.

104. Hugouvieux-Cotte-Pattat N, Condemine G, Shevchik VE. 2014. Bacterial pectate lyases, structural and functional diversity. Environ Microbiol Rep 6:427–440.

105. Jumper J, Evans R, Pritzel A, Green T, Figurnov M, Ronneberger O, Tunyasuvunakool K, Bates R, Žídek A, Potapenko A, Bridgland A, Meyer C, Kohl SAA, Ballard AJ, Cowie A, Romera-Paredes B, Nikolov S, Jain R, Adler J, Back T, Petersen S, Reiman D, Clancy E, Zielinski M, Steinegger M, Pacholska M, Berghammer T, Bodenstein S, Silver D, Vinyals O, Senior AW, Kavukcuoglu K, Kohli P, Hassabis D. 2021. Highly accurate protein structure prediction with AlphaFold. Nature 596:583–589.

106. Palanivelu P. 2006. Polygalacturonase: Active site analyses and mechanism of action. Indian J Biotechnol 5:148–162.

107. Herrera CM. 2017. Scavengers that fit beneath a microscope lens. Ecology 98:2725–2726.

108. Radja A, Horsley EM, Lavrentovich MO, Sweeney AM. 2019. Pollen Cell Wall Patterns Form from Modulated Phases. Cell 176:856–868.e10.

109. Bosch M, Hepler PK. 2005. Pectin Methylesterases and Pectin Dynamics in Pollen Tubes. Plant Cell 17:3219.

110. Engel P, Martinson VG, Moran NA. 2012. Functional diversity within the simple gut microbiota of the honey bee. Proc Natl Acad Sci U S A 109:11002–7.

111. Love RR, Weisenfeld NI, Jaffe DB, Besansky NJ, Neafsey DE. 2016. Evaluation of DISCOVAR de novo using a mosquito sample for cost-effective short-read genome assembly. BMC Genomics 17.

112. Parks DH, Imelfort M, Skennerton CT, Hugenholtz P, Tyson GW. 2015. CheckM: assessing the quality of microbial genomes recovered from isolates, single cells, and metagenomes. Genome Res 25:1043–1055.

113. Brettin T, Davis JJ, Disz T, Edwards RA, Gerdes S, Olsen GJ, Olson R, Overbeek R, Parrello B, Pusch GD, Shukla M, Thomason JA, Stevens R, Vonstein V, Wattam AR, Xia F. 2015. RASTtk: A modular and extensible implementation of the RAST algorithm for building custom annotation pipelines and annotating batches of genomes. Sci Rep 5:8365.

114. Asnicar F, Thomas AM, Beghini F, Mengoni C, Manara S, Manghi P, Zhu Q, Bolzan M, Cumbo F, May U, Sanders JG, Zolfo M, Kopylova E, Pasolli E, Knight R, Mirarab S, Huttenhower C, Segata N. 2020. Precise phylogenetic analysis of microbial isolates and genomes from metagenomes using PhyloPhlAn 3.0. Nature Communications 2020 11:1 11:1–10.

115. Katoh K, Standley DM. 2013. MAFFT Multiple Sequence Alignment Software Version 7: Improvements in Performance and Usability. Mol Biol Evol 30:772.

116. Nguyen LT, Schmidt HA, Von Haeseler A, Minh BQ. 2015. IQ-TREE: A Fast and Effective Stochastic Algorithm for Estimating Maximum-Likelihood Phylogenies. Mol Biol Evol 32:268.

117. Zharkikh A. 1994. Estimation of evolutionary distances between nucleotide sequences. J Mol Evol 39:315–329.

118. Csuos M. 2010. Count: evolutionary analysis of phylogenetic profiles with parsimony and likelihood. Bioinformatics 26:1910–1912.

119. Kluge AG, Farris JS. 1969. Quantitative Phyletics and the Evolution of Anurans. Syst Biol 18:1–32.

120. Syberg-Olsen MJ, Garber AI, Keeling PJ, McCutcheon JP, Husnik F. 2022. Pseudofinder: Detection of Pseudogenes in Prokaryotic Genomes. Mol Biol Evol 39.

121. Bateman A, Martin MJ, Orchard S, Magrane M, Agivetova R, Ahmad S, Alpi E, Bowler- Barnett EH, Britto R, Bursteinas B, Bye-A-Jee H, Coetzee R, Cukura A, da Silva A, Denny P, Dogan T, Ebenezer TG, Fan J, Castro LG, Garmiri P, Georghiou G, Gonzales L, Hatton-Ellis E, Hussein A, Ignatchenko A, Insana G, Ishtiaq R, Jokinen P, Joshi V, Jyothi D, Lock A, Lopez R, Luciani A, Luo J, Lussi Y, MacDougall A, Madeira F, Mahmoudy M, Menchi M, Mishra A, Moulang K, Nightingale A, Oliveira CS, Pundir S, Qi G, Raj S, Rice D, Lopez MR, Saidi R, Sampson J, Sawford T, Speretta E, Turner E, Tyagi N, Vasudev P, Volynkin V, Warner K, Watkins X, Zaru R, Zellner H, Bridge A, Poux S, Redaschi N, Aimo L, Argoud-Puy G, Auchincloss A, Axelsen K, Bansal P, Baratin D, Blatter MC, Bolleman J, Boutet E, Breuza L, Casals-Casas C, de Castro E, Echioukh KC, Coudert E, Cuche B, Doche M, Dornevil D, Estreicher A, Famiglietti ML, Feuermann M, Gasteiger E, Gehant S, Gerritsen V, Gos A, Gruaz-Gumowski N, Hinz U, Hulo C, Hyka- Nouspikel N, Jungo F, Keller G, Kerhornou A, Lara V, le Mercier P, Lieberherr D, Lombardot T, Martin X, Masson P, Morgat A, Neto TB, Paesano S, Pedruzzi I, Pilbout S, Pourcel L, Pozzato M, Pruess M, Rivoire C, Sigrist C, Sonesson K, Stutz A, Sundaram S, Tognolli M, Verbregue L, Wu CH, Arighi CN, Arminski L, Chen C, Chen Y, Garavelli JS, Huang H, Laiho K, McGarvey P, Natale DA, Ross K, Vinayaka CR, Wang Q, Wang Y, Yeh LS, Zhang J, Ruch P, Teodoro D. 2021. UniProt: the universal protein knowledgebase in 2021. Nucleic Acids Res 49:D480–D489.

122. Antipov D, Hartwick N, Shen M, Raiko M, Lapidus A, Pevzner PA. 2016. plasmidSPAdes: assembling plasmids from whole genome sequencing data. Bioinformatics 32:3380–3387.

123. Yang Z. 2007. PAML 4: phylogenetic analysis by maximum likelihood. Mol Biol Evol 24:1586–1591.

124. Zhang J, Nielsen R, Yang Z. 2005. Evaluation of an Improved Branch-Site Likelihood Method for Detecting Positive Selection at the Molecular Level. Mol Biol Evol 22:2472– 2479.

125. Mirdita M, Schütze K, Moriwaki Y, Heo L, Ovchinnikov S, Steinegger M. 2022. ColabFold: making protein folding accessible to all. Nat Methods 19:679–682.

126. Pettersen EF, Goddard TD, Huang CC, Meng EC, Couch GS, Croll TI, Morris JH, Ferrin TE. 2021. UCSF ChimeraX: Structure visualization for researchers, educators, and developers. Protein Sci 30:70–82.

127. Anandham R, Weon H-Y, Kim S-J, Kim Y-S, Kim B-Y, Kwon S-W. 2010. Acinetobacter brisouii sp. nov., isolated from a wetland in Korea. The Journal of Microbiology 48:36– 39.

128. Liu S, Wang Y, Ruan Z, Ma K, Wu B, Xu Y, Wang J, You Y, He M, Hu G. 2017. Acinetobacter larvae sp. nov., isolated from the larval gut of Omphisa fuscidentalis. Int J Syst Evol Microbiol 67:806–811.

129. Vaz-Moreira I, Novo A, Hantsis-Zacharov E, Lopes AR, Gomila M, Nunes OC, Manaia CM, Halpern M. 2011. Acinetobacter rudis sp. nov., isolated from raw milk and raw wastewater. Int J Syst Evol Microbiol 61:2837–2843.

