## Supplemental Figures for "Genome evolution following an ecological shift in nectar-dwelling *Acinetobacter*"

### Supplementary Figures

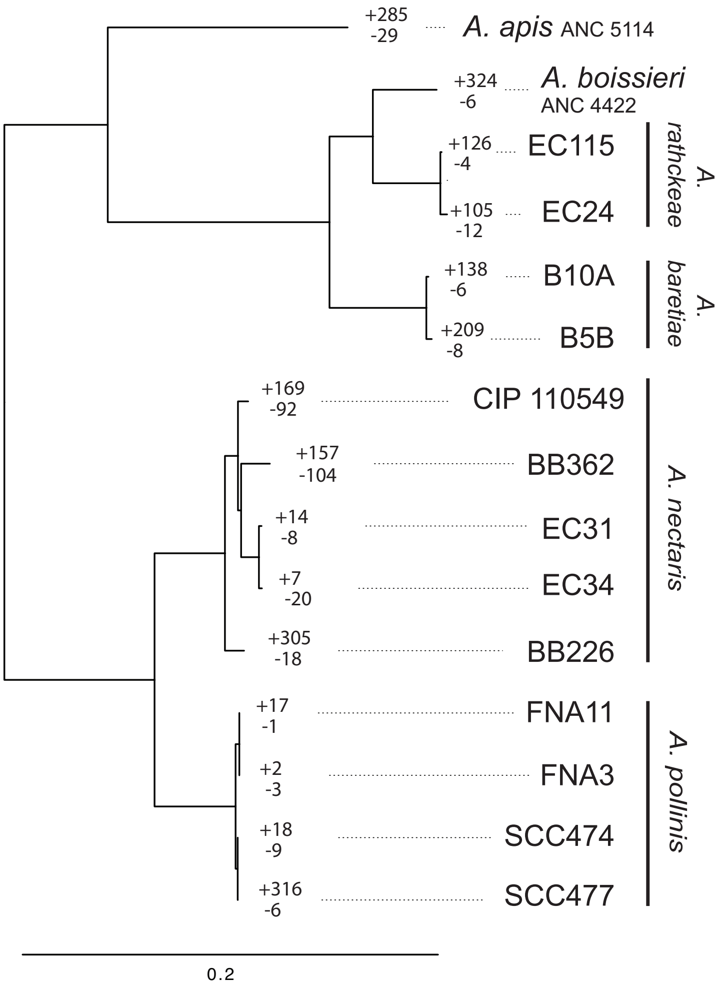


Figure S1: Gene gains and losses at tips within the nectar-dwelling clade. A subset of the phylogenomic tree from Fig. 1 is shown, with gains and losses at each tip given with a (+) or (-), respectively.

A

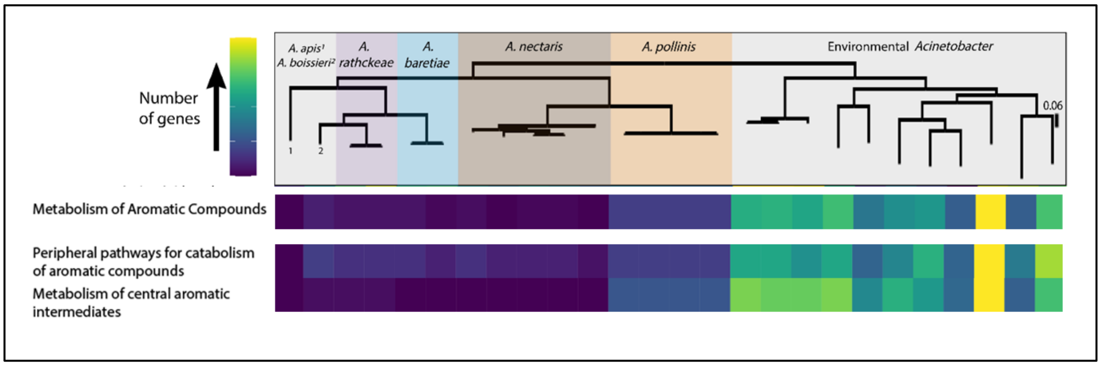


B

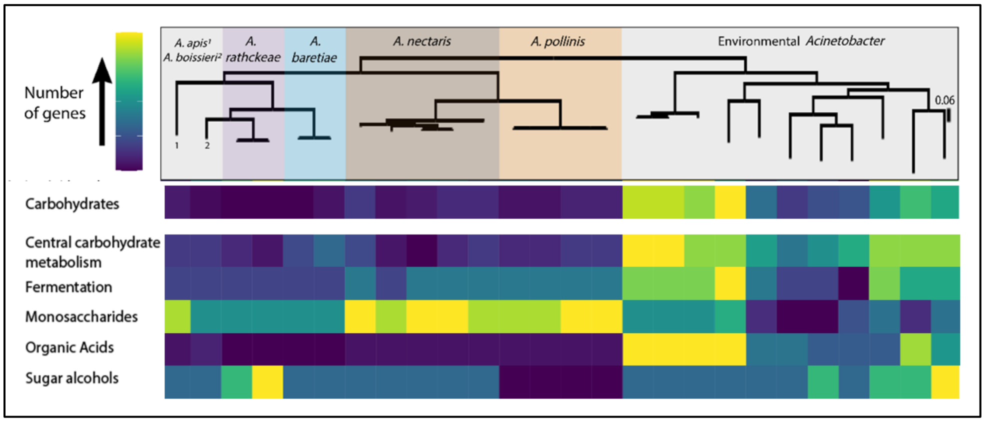


Figure S2: Heatmap showing number of orthologs across functional categories from nectar-dwelling and environmental *Acinetobacter* strains. Subcategory assignments expanded from Figure 1 are shown. A) RAST annotated subcatecories for the carbohydrate metabolism. B) RAST annotated subcategories for the metabolism of aromatic compounds. Each row is colored independently based on the range of gene content within the category. A color scale with yellow indicating the highest number of genes and deep blue indicating a lowest number of genes is used.

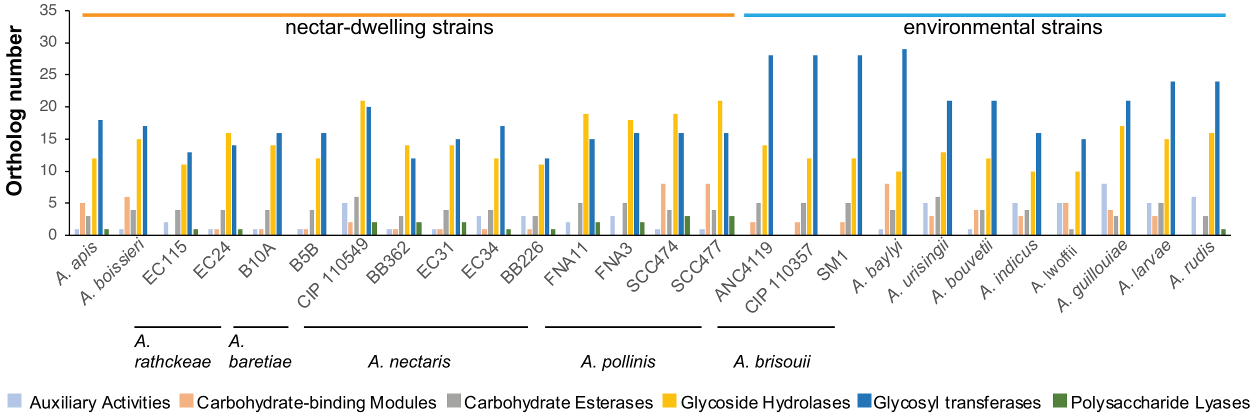


Figure S3: Number of orthologs in CAZy functional categories from environmental and nectar-dwelling *Acinetobacter* genomes.

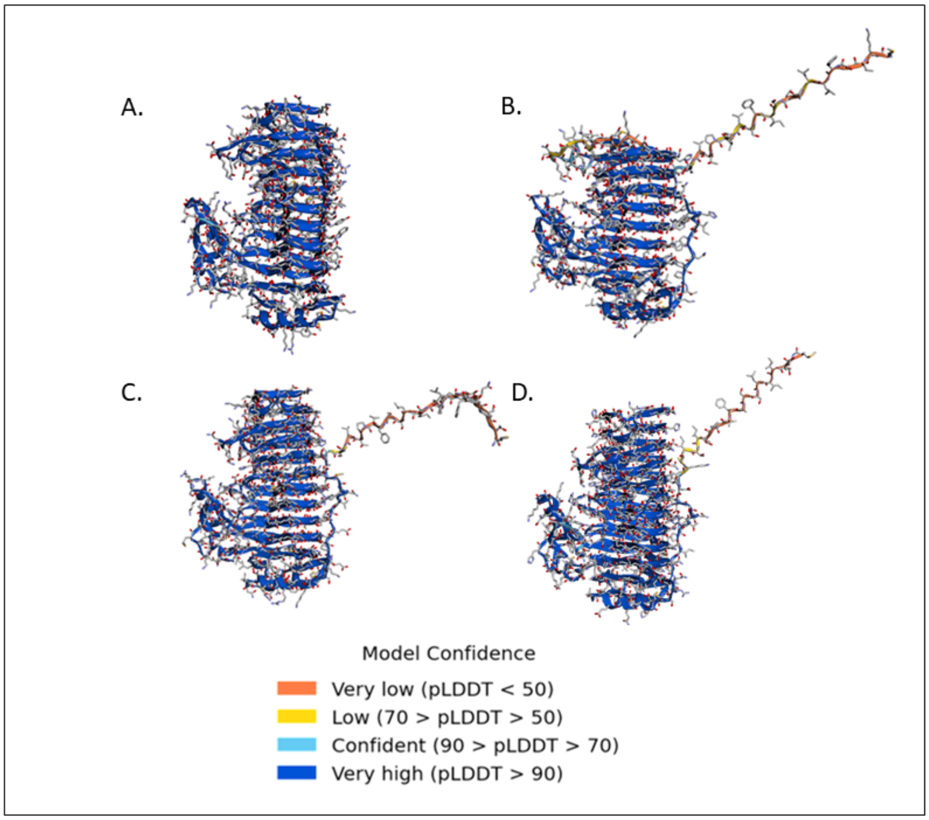


Figure S4: Alphafold protein predictions including confidence scores of polygalacturonase genes from *A. apis* (A), *A. pollinis* FNA3 (B-C), and *Phaseolibacter flectens* (D). The pLDDT scores are a per-residue estimate of confidence on a scale from 1-100. Protein structures are rotated here relative to Fig. 4 to show secretion signal tags clearly extending off the protein to the right. These portions of the proteins have low model confidence.

**Supplementary Tables**

Table S1: All orthologs from the clustering analysis with RAST functional categories shown. Orthologs involved in amino acid metabolism and transport are given in a separate sheet.

File name: Orthologs_functional_categories.xlsx

Table S1: Island viewer and Phaster results

| Strain | Island Count | Gene count on Islands | Number of intact prophage | Plasmids |
|---|---|---|---|---|
| EC31 | 7 | 111 | 1 | 7 |
| EC34 | 7 | 123 | 1 | 7 |
| BB362 | 8 | 189 | 2 | 10 |
| EC115 | 11 | 210 | 0 | 1 |
| EC24 | 11 | 188 | 3 | 0 |
| BB226 | 12 | 256 | 0 | 21 |
| B5B | 13 | 326 | 2 | 26 |
| B10A | 14 | 352 | 0 | 22 |
| FNA11 | 14 | 286 | 0 | 4 |
| FNA3 | 15 | 248 | 0 | 6 |
| SCC474 | 15 | 310 | 3 | 23 |
| SCC477 | 15 | 332 | 3 | 25 |

Table S3: Copies of pectin degrading genes within the nectar-dwelling *Acinetobacter* clade. Strains that contained at least one copy of pectin degrading genes in genomic islands are marked with an asterisk.

| Strain | Pectinlyase | Polygalacturonase |
|---|---|---|
| SCC474 | *3 | 4 |
| SCC477 | *3 | *6 |
| FNA11 | 2 | *5 |
| FNA3 | *2 | *5 |
| EC031 | 0 | 1 |
| EC034 | 1 | 1 |
| BB226 | 0 | 1 |
| BB362 | 1 | 1 |
| EC024 | 0 | 1 |
| EC115 | 0 | 1 |
| B10A | 0 | 1 |
| B5B | 0 | 1 |
| *A. boissieri* | 0 | 2 |
| *A. apis* | 1 | 1 |

Table S4: PAML amino acid selection test hits within polygalacturonase and pectin lyase orthologs.

File name: Selection_pectin_genes.xlsx
